# Novel metagenomics analysis of stony coral tissue loss disease

**DOI:** 10.1101/2024.01.02.573916

**Authors:** Jakob M. Heinz, Jennifer Lu, Lindsay K. Huebner, Steven L. Salzberg, Markus Sommer, Stephanie M. Rosales

## Abstract

Stony coral tissue loss disease (SCTLD) has devastated coral reefs off the coast of Florida and continues to spread throughout the Caribbean. Although a number of bacterial taxa have consistently been associated with SCTLD, no pathogen has been definitively implicated in the etiology of SCTLD. Previous studies have predominantly focused on the prokaryotic community through 16S rRNA sequencing of healthy and affected tissues. Here, we provide a different analytical approach by applying a bioinformatics pipeline to publicly available metagenomic sequencing samples of SCTLD lesions and healthy tissues from four stony coral species. To compensate for the lack of coral reference genomes, we used data from apparently healthy coral samples to approximate a host genome and healthy microbiome reference. These reads were then used as a reference to which we matched and removed reads from diseased lesion tissue samples, and the remaining reads associated only with disease lesions were taxonomically classified at the DNA and protein levels. For DNA classifications, we used a pathogen identification protocol originally designed to identify pathogens in human tissue samples, and for protein classifications, we used a fast protein sequence aligner. To assess the utility of our pipeline, a species-level analysis of a candidate genus, *Vibrio*, was used to demonstrate the pipeline’s effectiveness. Our approach revealed both complementary and unique coral microbiome members compared to a prior metagenome analysis of the same dataset.

**Article Summary:** Studies of stony coral tissue loss disease (SCTLD), a devastating coral disease, have primarily used 16S rRNA sequencing approaches to identify putative pathogens. In contrast, this study applied human tissue pathogen identification protocols to SCTLD metagenomic DNA samples. Diseased samples were filtered of host sequences using a k-mer based method since host genomes were unavailable. DNA and protein-level classifications from this novel approach revealed both complementary and unique microbiome members compared to a prior metagenome analysis of the same dataset.

## Introduction

Stony coral tissue loss disease (SCTLD) was discovered off the coast of Miami, FL in 2014 and has since had negative consequences on the function of coral reefs across Florida and the Caribbean (Walton *et al*. 2018; Alvarez-Filip *et al*. 2022). To date, despite many efforts, no pathogen has been definitively identified as the causative agent of SCTLD. The stony coral (order Scleractinia) microbiome is a complex system of interactions between the host, bacteria, viruses, fungi, archaea, and algal symbionts (Bourne *et al*. 2009); thus a disturbance in any number of these symbiotic relationships could be involved in SCTLD progression. Multiple studies have explored viruses that may infect stony coral symbionts, notably *Symbiodiniaceae*, but no causative relationships have been detected (Work *et al*. 2021; Veglia *et al*. 2022; Beavers *et al*. 2023; Howe-Kerr *et al*. 2023). Bacterial species are particularly under scrutiny for their potential involvement in SCTLD, due to the effectiveness of antibiotics in halting lesion progression in multiple affected coral species (Aeby *et al*. 2019, Neely *et al*. 2020; Shilling *et al*. 2021; Studivan *et al*. 2023). Consequently, SCTLD studies have predominantly focused on understanding changes in the bacterial community between apparently healthy and SCTLD-affected corals.

Studies to identify bacterial pathogens have relied primarily on small subunit 16S ribosomal RNA (rRNA) sequencing, followed by computational analysis (Callahan *et al*. 2016) typically using the Silva database (Quast *et al*. 2013) to assign and classify Amplicon Sequence Variants (ASVs) into taxa. ASVs found in diseased lesion samples are then compared to samples from apparently healthy colonies to determine which ASVs are associated with the tissue loss lesions (Meyer *et al*. 2019; Rosales *et al*. 2020; Clark *et al*. 2021). These methods have characterized many notable shifts in coral bacterial communities due to SCTLD and several bacterial taxa associated with SCTLD lesions, including *Rhizobiales*, *Clostridiales*, *Peptostreptococcales-Tissierellales*, *Rhodobacteraceae*, *Flavobacteriaceae*, and *Vibrionaceae* (Rosales *et al*. 2023). However, because of the difficulty in determining whether an associated bacterial taxon is a harmless commensal, an opportunistic secondary infection, or the primary pathogen, none of the bacterial taxa associated with SCTLD have been identified as the causative agent.

An alternative approach to understanding disease dynamics is the use of metagenomic sequencing, in which all of the DNA from a source is sequenced, including not only the host, but also viruses, bacteria, and eukaryotic species living on or within the host tissue. For example, by analyzing metagenomic sequencing data of human tissue samples taken from the site of infection, researchers have identified pathogenic agents in brain infections (Salzberg *et al*. 2016; Wilson *et al*. 2019), corneal infections (Eberhart *et al*. 2017), and other diseases (Kostic *et al*. 2012). The sensitivity of this approach relies on first, sequencing the source DNA deeply enough to capture the pathogen of interest, and second, the existence of genome assemblies closely related to the pathogen in public databases. While the number of complete genomes has grown enormously over the past two decades, databases still contain few or no genomes for non-model organisms, including scleractinian corals.

Currently, data from only one SCTLD metagenome study is publicly available. While the authors of that study (Rosales *et al*. 2022) were able to assemble and annotate genomes for SCTLD-associated bacterial taxa such as *Rhodobacterales*, *Rhizobiales*, and *Flavobacteriales*, the results were focused on only five of the twenty diseased lesion tissue samples, all from the same coral species (*Stephanocoenia intersepta*), because the majority of samples were dominated by host sequences. In metagenomic studies, host sequences can confound results, so they are typically removed by aligning all reads to a host reference genome (Gihawi *et al*. 2023; Lu *et al*. 2022). Currently, the GenBank database has 53 genome assemblies from scleractinian corals, of which only seven are at the chromosome level (NCBI 2023). Of these 53 genomes, none are from the species of corals previously investigated for SCTLD (Rosales *et al*. 2022), emphasizing the additional challenges associated with using metagenomics in non-model organisms. Additionally, given the complex symbiotic microbiome (i.e., algal symbiont, viruses, and prokaryotic community) of stony corals (Bourne *et al*. 2009), the host DNA is only one of the hurdles.

In this study, we applied new classification methods to understand this devastating coral disease. We used a method to filter host reads from metagenome data by using data collected from apparently healthy corals of the same species to approximate a species-specific healthy host coral genome and microbiome. We then applied the Kraken software suite for pathogen identification (Lu *et al*. 2022) using KrakenUniq (Breitwieser *et al*. 2018) to identify putative pathogens present in diseased samples and not present in healthy ones. Using these methods, we identified a number of taxa that have previously been associated with SCTLD, providing further support for their involvement in SCTLD pathogenesis. Finally, from the pool of bacterial taxa we identified as associated with SCTLD lesions, we selected a candidate genus, *Vibrio*, with which to explore the utility of our pipeline at a finer taxonomic level.

## Materials and methods

### Data acquisition

We downloaded 58 metagenomic datasets that consisted of 150 bp paired-end reads from NCBI Bioproject PRJNA576217 (Benson *et al*. 2017), previously generated by Rosales *et al*. (2020). Sample SRR15960000, an apparently healthy *Diploria labyrinthiformis* sample, was removed due to data quality problems, leaving 57 sets of paired-end samples for analysis. These were 20 diseased colony lesion (DL) samples, 20 diseased colony unaffected (DU) samples, and 17 apparently healthy colony (AH) samples from the coral species *D. labyrinthiformis*, *Dichocoenia stokesii*, *Meandrina meandrites*, and *Stephanocoenia intersepta*. DU samples were taken from apparently unaffected tissue from the diseased corals also sampled for DL. All samples were collected and treated in the same manner, as previously described (Rosales *et al*. 2020, 2022). In brief, coral tissue and mucus slurries were scraped from the coral surface with 10-ml plastic syringes, transferred to plastic tubes on the boat, and held in a cooler on ice until being flash-frozen in a liquid nitrogen dewar on shore; samples were held in a −80℃ freezer until processed for sequencing. Because all samples were from corals within reefs with an ongoing SCTLD outbreak in the Florida Keys, it is possible that a primary pathogen of SCTLD could be present in low abundance in at least one of the AH samples or that the AH microbiome was different from that of corals in reefs where SCTLD had yet to arrive. Therefore, in this study, our findings represent microbial communities associated with the observable surface tissue loss formation stage of SCTLD (hereafter visual tissue loss or diseased lesion) compared to corals with no visual signs of disease (i.e., AH).

The tissue samples from the four coral species were pooled by each of the three disease states, resulting in twelve pooled read files (AH, DU, and DL for each of the four coral species). It was assumed that a putative pathogen involved in visual tissue loss would likely show different abundances in DL samples during different stages of lesion progression, so pooling the samples was thought to increase the likelihood of observing a putative agent. All subsequent analyses were based on these data, focusing primarily on the DL and AH samples. The reads from the DU samples were explored in the preliminary analysis but were not considered in the final analysis. Due to the proximity of the DU samples to lesion tissue, DU samples were considered likely to represent early stages of surface tissue loss, and therefore poor choices for our methods.

### Filtering reads with a healthy coral reference database

Because no sequenced genome was available for any of the four coral species, we created a customized database to identify reads that likely originated from either the host genome or the healthy host microbiome. To do this, we used reads from all AH samples to create a KrakenUniq (Breitwieser *et al*. 2018) database for each coral species. We then used this database along with KrakenUniq to classify reads from DL samples, thereby removing any read in DL samples that matched any read in the AH samples. This filtering step produced a subset of diseased reads that we considered unique to the DL samples, and greatly reduced the number of reads analyzed in subsequent steps (Figure 1a).

**Figure 1:**
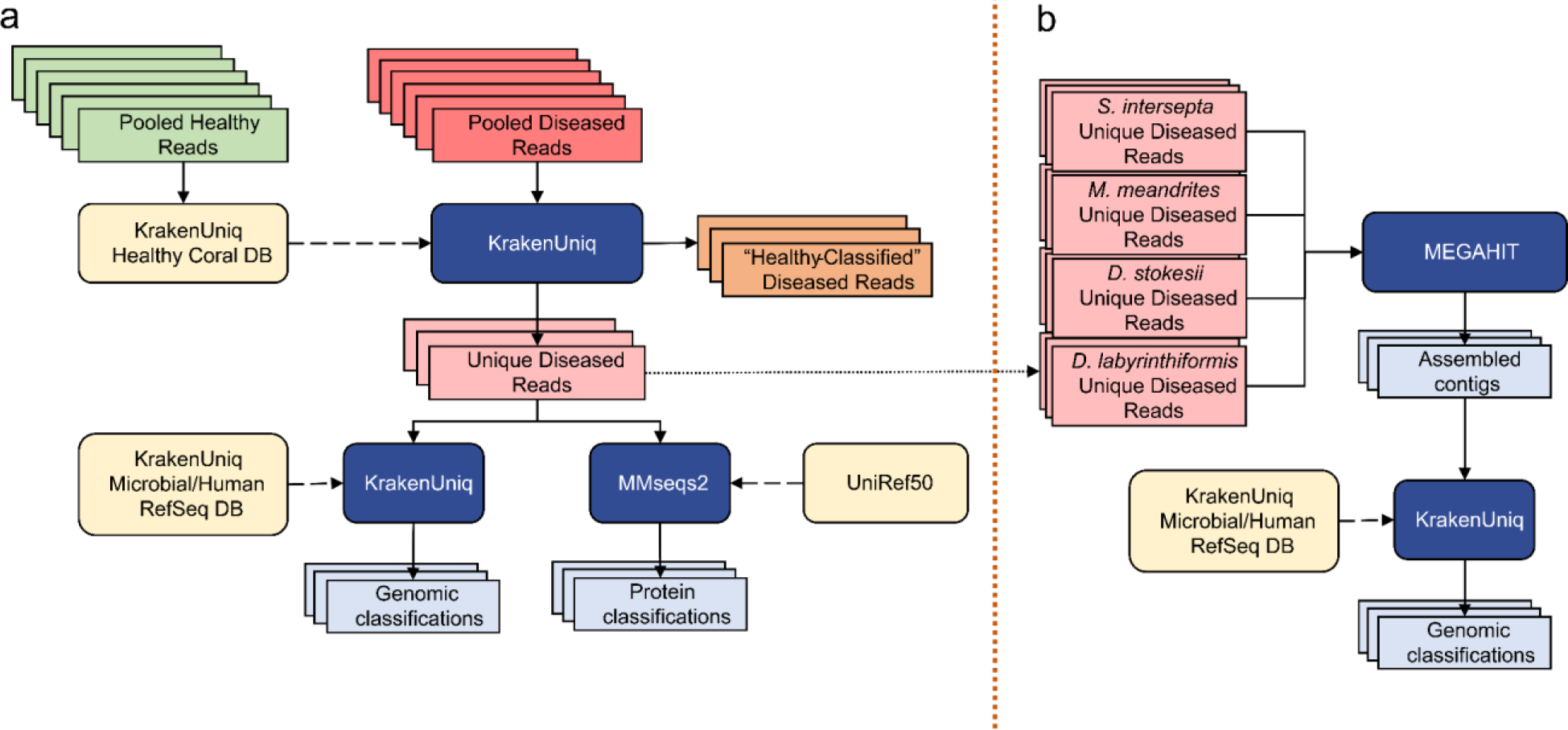
**(a)** Filtering the diseased reads consisted of building a KrakenUniq database from all healthy reads for each coral species and classifying the corresponding pooled diseased reads against their respective database. That is every coral species was only filtered with its respective “Healthy” Coral DB, and there was no cross-species filtration. Reads that were unclassified by the database were considered unique to the diseased samples. This subset was classified at the DNA level with KrakenUniq and at the protein level with MMseqs2. **(b)** The unique diseased reads from all coral species were combined and assembled with MEGAHIT. The assembled contigs were then classified with KrakenUniq.

The k-mer size for databases was set to 29 bp, lower than the default of 31bp, because we wanted to filter more aggressively. For all other parameters, the default values of KrakenUniq were used. For each coral species, the pooled DL reads were classified with KrakenUniq against the AH reads database corresponding to that species. The original DL files were parsed to extract all reads that were unclassified by the AH KrakenUniq database, generating the DL unique files used for subsequent analysis (Figure 1a). With this aggressive filtration approach, we may have lost information about changes in relative abundances between the AH and DL samples and were therefore limited to only investigating species in high abundance, but we were left with reads that were truly unique to the DL samples. This set would likely provide the clearest signal of microbes associated with visual lesion formation, which was the primary goal of this study.

### Genus-level classification with KrakenUniq

To first establish a baseline abundance profile before any filtering, the AH and DL reads were classified with KrakenUniq (Breitwieser *et al*. 2018) using default parameters and the default k-mer size of 31 bps against a microbial database (Figure 1a). The database used for classification was built in August 2020 using all NCBI RefSeq complete bacterial, viral, and archaeal genomes, the GRCh38 human genome, the NCBI UniVec database, and a curated set of sequences from EuPathDB (Lu and Salzberg 2018; Amos *et al*. 2022). Then, the unique DL reads were also classified with KrakenUniq using the same database and parameters. All reads that were classified as *Homo* (human contamination) by KrakenUniq were removed from subsequent analysis and results. If there are novel species associated with SCTLD, then their genomes will not be present in public databases; however, if closely related species from the same genera are available, then we might find DNA sequence-level matches to those genomes. For this reason, the unique k-mers are reported at the genus level. Additionally, the relative abundances of the genera are calculated by the unique k-mer count rather than the read count. In general, using the unique k-mer count (i.e., sequences of length k are counted just once per taxon, no matter how many times they occur in the raw data) rather than read count reduces the bias introduced from using amplification-based sequencing workflows. Using the unique k-mer count also reduces false positives that may arise from reads that contain low-complexity k-mers (Breitwieser *et al*. 2018).

The report files were initially visualized and explored with Pavian (Breitwieser and Salzberg 2020). The read classifications were verified by randomly sampling 40 classified reads, aligning them with megablast (Altschul *et al*. 1990) to the NCBI standard non-redundant (nr) nucleotide database, and ensuring they had the same or similar classifications as with KrakenUniq. The unique k-mers-per-read statistic served as a confidence flag. For a species that was truly present in the sample, even with amplified metagenomic data, we expected a high number of unique k-mers per read. A 150 bp read may contain up to 120 unique 31-mers, although repetitive k-mers will reduce the unique count. There is also an upper bound to unique k-mers found in a genome, which may be reached when the genome is small or when the sampling depth is high. In this study, we considered a value of less than five unique k-mers per read as a flag that the taxon might be a false positive. The unique k-mer-per-read count for every classified genus is reported in Supp. Table S1.

### Protein-level classification with MMseqs2

Because protein sequences are more conserved than DNA across distant species, we ran translated searches using MMSeqs2 easy-taxonomy workflow (Steinegger and Söding 2017) with the UniRef50 protein database (Suzek *et al*. 2015) to determine if this would identify more of the microbial reads than DNA sequences alone (Figure 1a). UniRef50 (Suzek *et al*. 2015) allowed for faster alignment than using the standard UniRef given that we aligned our protein sequences to clusters of similar protein sequences rather than all protein sequences. The paired reads had to be classified separately because MMseqs2 easy-taxonomy does not allow for both paired reads to be processed together. Additionally, MMSeqs2 does not report k-mer counts, only read counts, a metric that is subject to more bias from PCR amplification protocols.

To identify reads belonging to members of the algal symbiont family *Symbiodiniaceae*, which were among the most abundant reads (see Results), the MMseqs2 output was parsed to extract all protein cluster identifiers that had at least one alignment at the “f_Symbiodiniaceae” level or below for each coral species. The UniRef50 cluster identifiers were mapped to the full UniProtKB (UniProt Consortium, 2021). Their functions, if known, are reported in Supp. Table S2 as output by the UniProt ID mapping service (Huang *et al*. 2011). Supp. Table S2 was produced by inserting the number of alignments from the original MMseqs2 “tophit_report” files into the outputs of the UniProtKB ID mapping.

### Contig assembly and classification

Due to the high genomic diversity of viruses (Aiewsakun *et al*. 2018), a viral agent might have been missed in our previous read analyses because it was too divergent from available DNA and protein sequences. This problem could be mitigated if the query sequences were longer, and therefore we assembled the raw reads to see if any long viral contigs were assembled. With our filtering approach, we did not expect to have the depth of coverage to assemble anything much larger than an abundant virus.

The filtered unique diseased reads from all four coral species were pooled to form a fasta file of all filtered unique diseased reads and were then assembled with MEGAHIT v1.2.9 (Li *et al*. 2016) using default parameters (Figure 1b). The contigs were classified with KrakenUniq using the same database of complete bacterial and viral genomes used above. The viral classifications from the report file were extracted to search for any viruses of interest. These steps were repeated with pooling just the filtered unique diseased reads excluding *S. intersepta,* because samples from this species represented a majority of the reads (89.8%) in our study and dominated the previous assembly (Rosales *et al*. 2022).

### Candidate genus (*Vibrio*) investigation

To explore the utility of our pipeline in characterizing the SCTLD microbiome at a finer taxonomic level, we selected a candidate bacterial genus, *Vibrio*, from the k-mers found most abundant by KrakenUniq (see Results). We selected *Vibrio* for this exercise because this genus was not emphasized in the previous metagenomic sequencing analysis of this same dataset (Rosales *et al*. 2022), *V. coralliilyticus* coinfection increases the virulence of SCTLD (Ushijima *et al*. 2020), and many *Vibrio* species are pathogenic to marine organisms broadly, not just to corals (de Souza Valente and Wan 2021; Dincturk *et al*. 2023). First, the reads that were classified to the *Vibrio* species level were investigated in further detail. To do this, the KrakenUniq report files of the unique diseased reads for every coral species were parsed to extract the number of unique k-mers assigned to each *Vibrio* species. The k-mer counts were normalized by dividing them by the total number of k-mers assigned to the *Vibrio* genus within the respective coral species. To investigate the prevalence of these *Vibrio* species across the samples comprising coral species data, we calculated the relative proportion of different *Vibrio* species for individual samples. The contribution of *Vibrio* species reads from each sample was then found by parsing the KrakenUniq output to determine the sample ID number from the read identifier. With these methods, it was not possible to determine the number of unique k-mers that were contributed by each sample, therefore the read counts were reported. Second, to compare our identified *Vibrio* reads with previous SCTLD *Vibrio* assemblies, the reads that were classified at or below the *Vibrio* genus level by KrakenUniq were extracted from the original sequence files for *S. intersepta* and *D. labyrinthiformis* only, because they contributed the majority of the *Vibrio* genus reads (see Results). The draft genomes from an SCTLD study that cultured *V. coralliilyticus* (Ushijima *et al*. 2020) were downloaded from NCBI Bioproject PRJNA625269 (Benson *et al*. 2017) and a Bowtie2 index was built for each one. The extracted *Vibrio* reads were aligned with Bowtie2 (Langmead and Salzberg 2012) to each of the eight draft genomes. The Bowtie2 alignment rates to each draft genome are reported in Supp. Table S4.

## Results

### Filtering reads with a healthy coral reference database

We analyzed metagenomic reads from corals *D. labyrinthiformis, D. stokesii, M. meandrites,* and *S. intersepta* from three different tissue sample types: diseased colony lesion (DL), diseased colony unaffected (DU), and apparently healthy colony (AH). The total number of reads from each species and sample type is shown in Table 1.

**Table 1:**
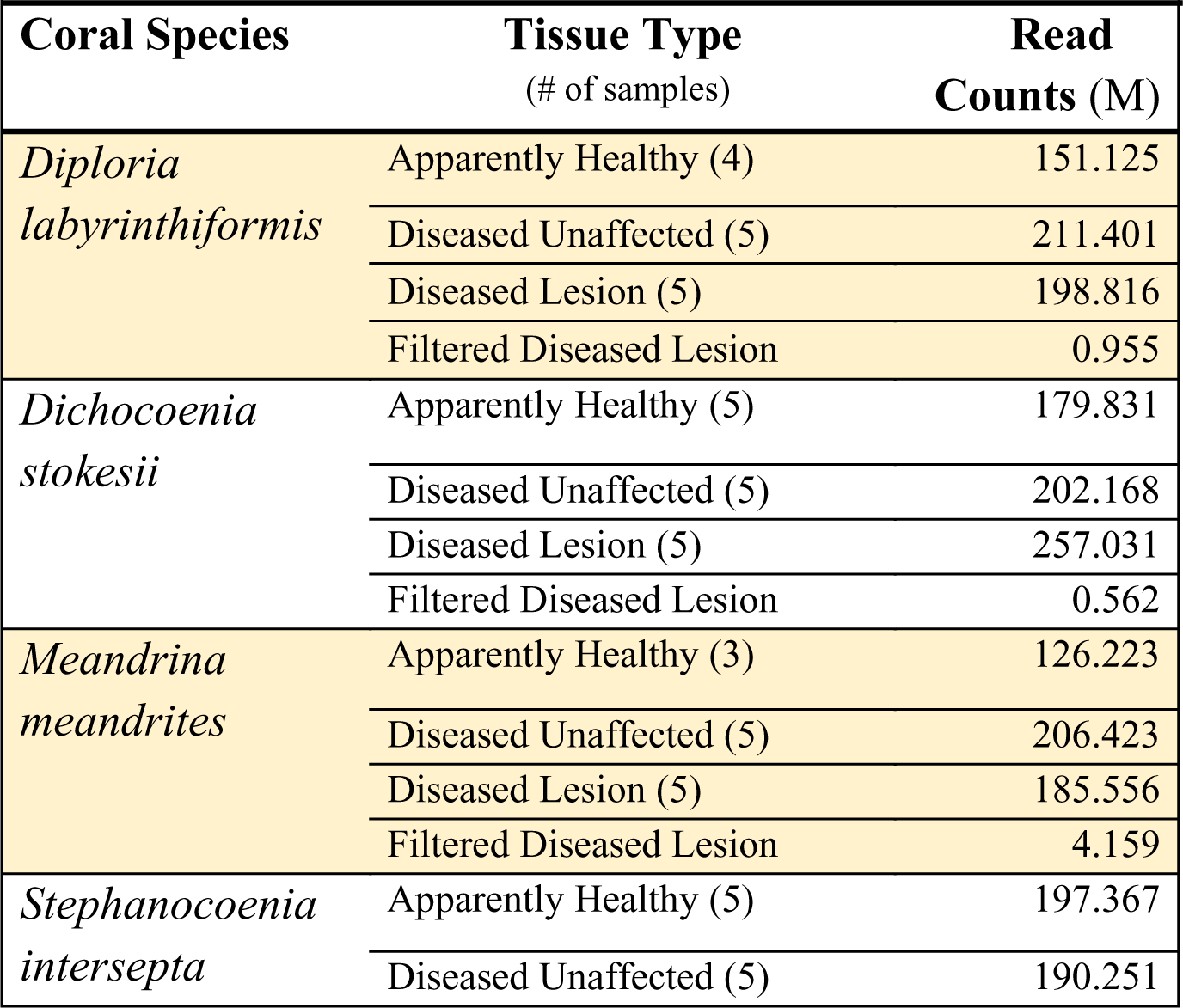
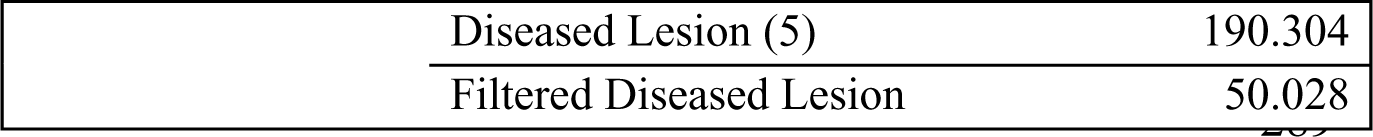
Summary of total DNA sequencing reads from each coral species and tissue type. Rows labeled “Filtered” report the unique reads remaining after filtering out reads that overlapped with those found in apparently healthy samples, as described in the text. M=millions of reads.

|  |  |  |
| --- | --- | --- |
| **Coral Species** | **Tissue Type**<br>(# of samples) | **Read Counts (M)** |
| *Diploria labyrinthiformis* | Apparently Healthy (4) | 151.125 |
|  | Diseased Unaffected (5) | 211.401 |
|  | Diseased Lesion (5) | 198.816 |
|  | Filtered Diseased Lesion | 0.955 |
| *Dichocoenia stokesii* | Apparently Healthy (5) | 179.831 |
|  | Diseased Unaffected (5) | 202.168 |
|  | Diseased Lesion (5) | 257.031 |
|  | Filtered Diseased Lesion | 0.562 |
| *Meandrina meandrites* | Apparently Healthy (3) | 126.223 |
|  | Diseased Unaffected (5) | 206.423 |
|  | Diseased Lesion (5) | 185.556 |
|  | Filtered Diseased Lesion | 4.159 |
| *Stephanocoenia intersepta* | Apparently Healthy (5) | 197.367 |
|  | Diseased Unaffected (5) | 190.251 |

|  |  |
| --- | --- |
| Diseased Lesion (5) | 190.304 |
| Filtered Diseased Lesion | 50.028 |

### Genus-level classification with KrakenUniq

Before any filtering, both the AH and the DL reads were profiled to establish baseline abundances. The AH reads were dominated by *Synechococcus* (Supp. Fig S1). Although filtering removed a significant majority (up to 99%) of the reads, the original DL microbial read classifications did not have notable changes (Supp. Fig S2), indicating predominantly coral host DNA was filtered from the DL samples, which led to substantial improvements in computational speed in downstream analyses.

When the KrakenUniq reports for filtered DL samples were sorted by unique k-mer count within coral species, the microbial genera *Synechococcus, Vibrio, Ruegeria, Phaeobacter,* and *Sulfitobacter* were found at high proportions in all coral species, with the genera *Synechococcus*, *Vibrio,* and *Ruegeria* being particularly abundant across all coral species (Figure 2). *Synechococcus* was the most or second most abundant in all coral species. In *D. labyrinthiformis, Vibrio* was the most abundant, with 4.1 times more unique k-mers than the second most abundant genus, *Synechococcus*. In *D. stokesii*, *Synechococcus* was the most abundant, with *Vibrio* being second most abundant. As in D. stokesii, *Synechococcus* was the most abundant in *M. meandrites*, but in this coral *Pseudovibrio* was the second most abundant and was the only coral in which this genus appeared in a high proportion. *Vibrio* had the sixth highest relative abundance in *M. meandrites,* which, though lower than observed in the other coral species, still represented a high k-mer-to-read ratio of 52.0 (Supp. Table S1). In *S. intersepta, Vibrio* was again clearly the most abundant, having 4.3 times the amount of unique k-mers compared to the second place *Synechococcus.* Due to their high abundances across all coral species in this analysis, *Synechococcus*, *Vibrio,* and the *Rhodobacteraceae* family (to which *Ruegeria, Phaeobacter,* and *Sulfitobacter* belong) appear to be associated with visual tissue loss and may represent important agents of SCTLD. Another useful aspect of the KrakenUniq pipeline is that by including the human reference genome in the database, human contamination can be detected in the classification step, without the expensive additional step of aligning all reads to the human reference genome. Human contamination, as reported by KrakenUniq, represented less than 0.2% of the filtered DL reads for all coral species. In the initial unfiltered DL and AH samples, human contamination was reported to be between 1.5 - 3.0% of reads across samples, an expected level for marine samples (Schmieder and Edwards 2011).

**Figure 2:**
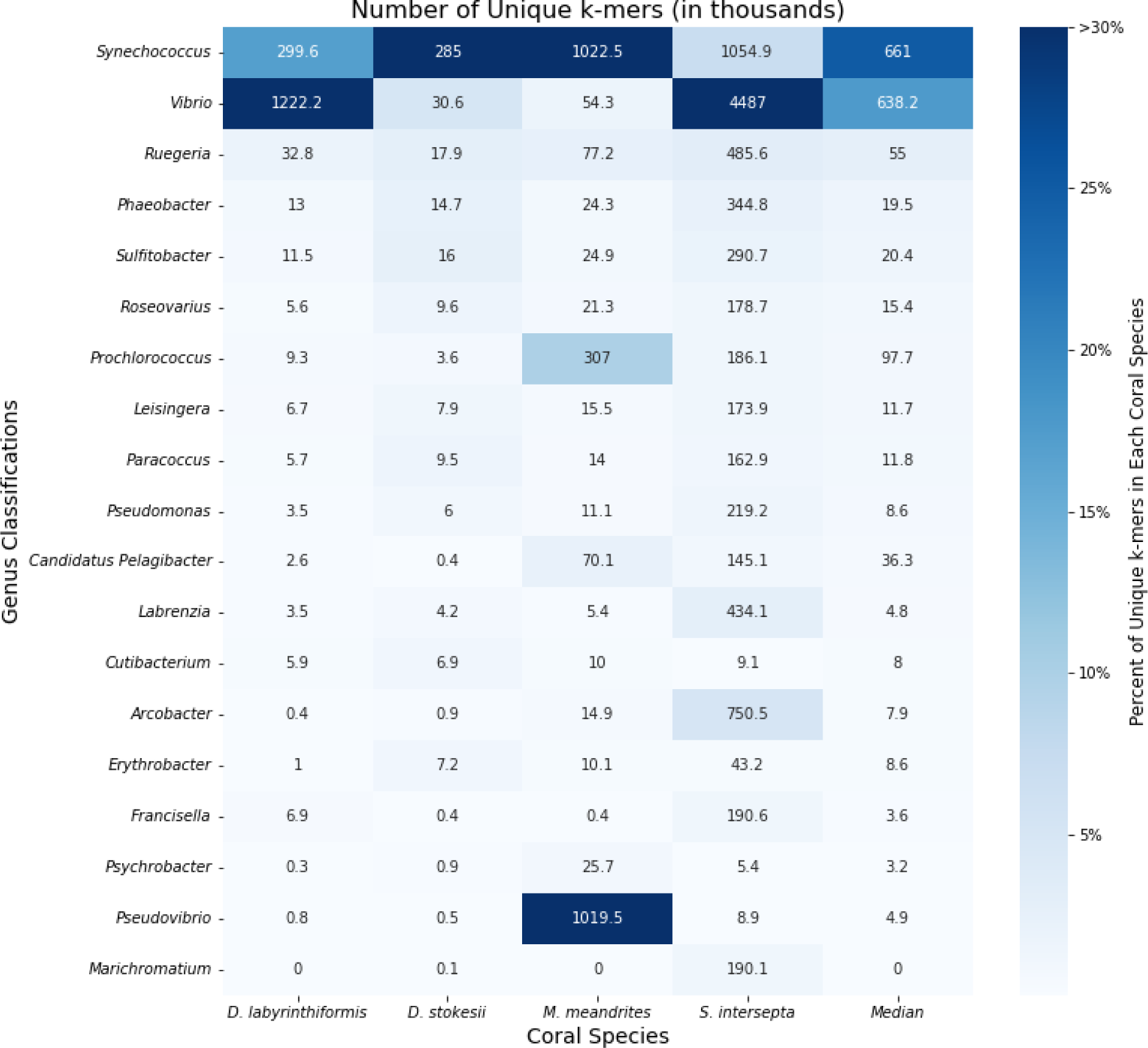
Unique k-mer counts from KrakenUniq genus-level classifications of unique diseased reads for every coral species. The intensity of the shading represents the percent of total unique k-mers assigned to the genus. Each box is annotated with the number of unique k-mers (in thousands) assigned to the genus.

### Protein-level classification with MMseqs2

The number of microbial reads classified for each coral species at the protein-level using MMseqs2 (Steinegger and Söding 2017) increased approximately six-fold compared to the DNA-based searches (Supp. Figure S3). Because we were primarily interested in whether any new candidate taxa emerged, we did not consider the relative abundances of different taxa classified by the protein-based search compared to the DNA-based search. Due to the decreased specificity of a protein search, MMseqs2 classified many reads as “unclassified [family level]”; thus, the results are presented at the family rather than the genus level (Figure 3). As expected, the read counts for each coral species were similar between the paired reads, which are classified separately by MMSeqs2.

**Figure 3.**
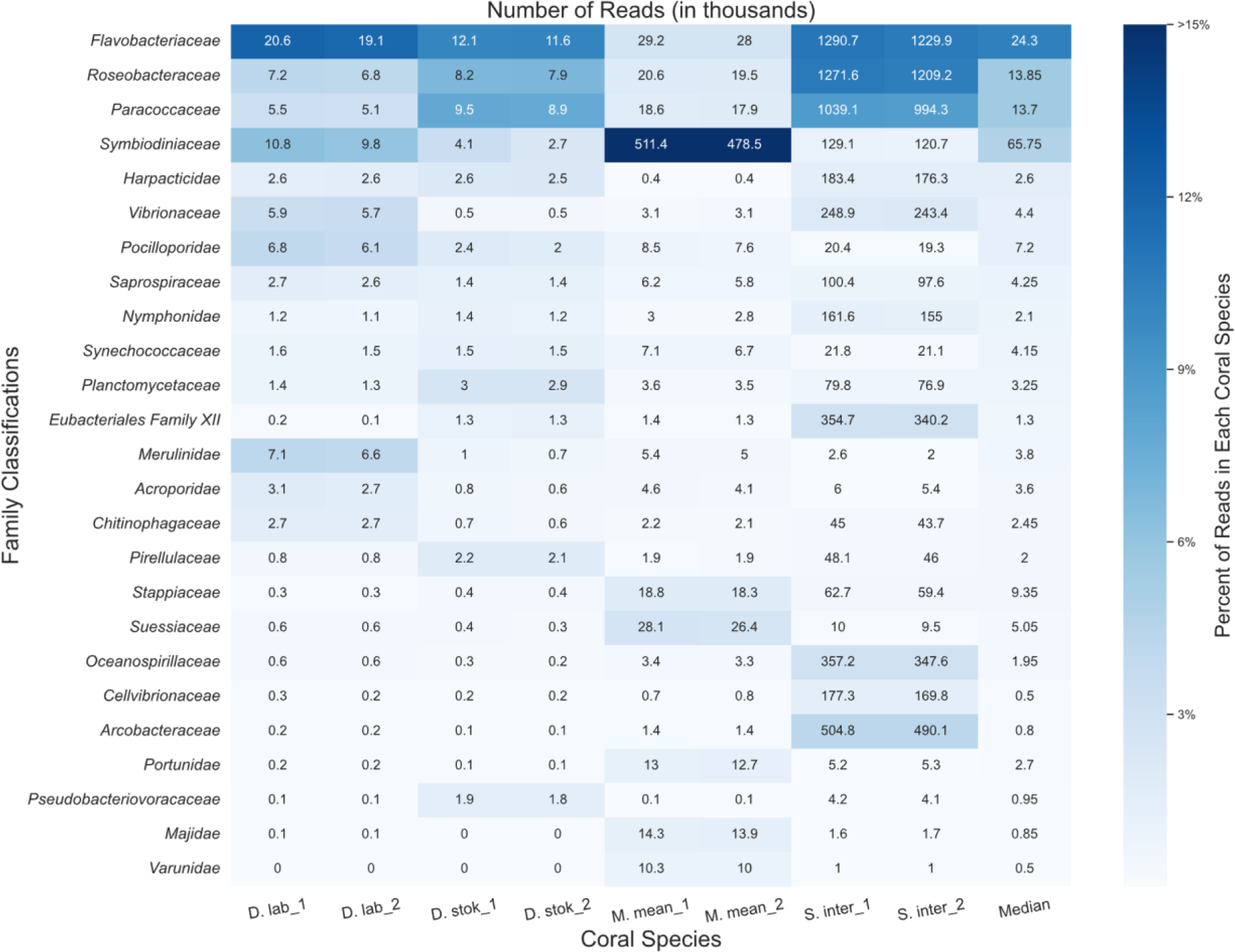
Classifications from MMseqs2 at the family-level for forward (“1”) and reverse reads (“2”) from unique diseased reads from each coral species. The intensity of the shading represents the proportion of total reads from the coral species that were assigned to the family. The boxes are annotated with the number of reads (in thousands) assigned to each family.

MMseqs2 was able to classify a higher percentage of bacterial and eukaryotic reads compared to the KrakenUniq DNA-level analysis (Supp. Figure 3), predominantly from coral algal symbionts such as *Symbiodiniaceae*, yet we saw similar abundant taxa. *Symbiodiniaceae* was among the top families in all coral species and particularly abundant (∼48% of all classified reads) in *M. meandrites*. Due to particular interest in the role of *Symbiodiniaceae* in SCTLD progression (Landsberg *et al*. 2020, Beavers *et al*. 2023), the functions of the proteins in the *Symbiodiniaceae* protein clusters identified are provided in Supp. Table S2.

The families *Flavobacteriaceae, Roseobacteraceae*, and *Paracoccaceae* were among the most abundant families in all coral species. In this database, *Roseobacteraceae* and *Paracoccaceae* are homotypic synonyms of the family *Rhodobacteraceae* in the database used for DNA-level classifications (Göker 2022). So, we observed *Rhodobacteraceae* as before in the DNA analysis, but *Flavobacteriaceae* emerged as another family of interest in this protein-level analysis.

*Flavobacteriaceae* was also found as a top family in the DNA-level classification (Supp. Figure S4); however, no top genus was identified that belongs to this family. Additionally, the average number of unique k-mers stemming from reads classified at or below the *Flavobacteriaceae* family at the DNA-level was relatively low. For example, in *S. intersepta*, there were 794,258 unique k-mers from 489,794 reads, or ∼1.6 k-mers per read. In *D. stokesii* there were 38,333 unique k-mers from 3,802 reads, or ∼10 k-mers per read. Across all coral species, the *Flavobacteriaceae* family had one of the lowest average k-mer-per-read counts of all bacterial families identified. For example, the *Vibrionaceae* family, which has similar sized genomes to *Flavobacteriaceae* (Lin *et al*. 2018; Gavriilidou *et al*. 2020), had 25 k-mers-per-read and 88 k-mers-per-read in *S. intersepta* and *D. stokesii*, respectively.

### Contig assembly and classification

In addition to characterizing the bacterial community, using metagenomic data made it possible to explore DNA viruses found in the unique DL reads. To account for the genomic diversity of viruses, which may not share many conserved sequences with genomes in public databases, we assembled contigs from the unique DL reads and classified them with KrakenUniq to identify any viral contigs that may be of interest in SCTLD etiology. This resulted in 1,014,402 assembled contigs. KrakenUniq classified 168,829 (16.6%) contigs, of which only 227 (0.02%) were viruses. *Paracoccus* phages were the most abundant viral contigs, with *Synechococcus* phages*, Dinoroseobacter* phages*, Vibrio* phages, and *Cyanophages* being abundant as well (Figure 4). These phage results reflect bacteria that were found in high proportions in the results above, providing support that the high abundances of the associated bacteria previously observed were representative of the true metagenomic compositions of the samples. Finally, five contigs were classified as *Chrysochromulina ericina* virus, a virus that infects the microalga *Chrysochromulina ericina* (also known as *Haptolina ericina*) (Gallot-Lavallée *et al*. 2017), but they represented only 108 unique k-mers and an assembly of 2.5 kb of a 473.6 kb genome (Gallot-Lavallée *et al*. 2015). When aligned with BLASTN (Altschul et al. 1990) to standard databases, only two contigs aligned best to *C. ericina* virus and the other three aligned best to Eukarya, possibly indicating false positives and leading us to be skeptical of any significant implications of this finding for SCTLD progression.

**Figure 4:**
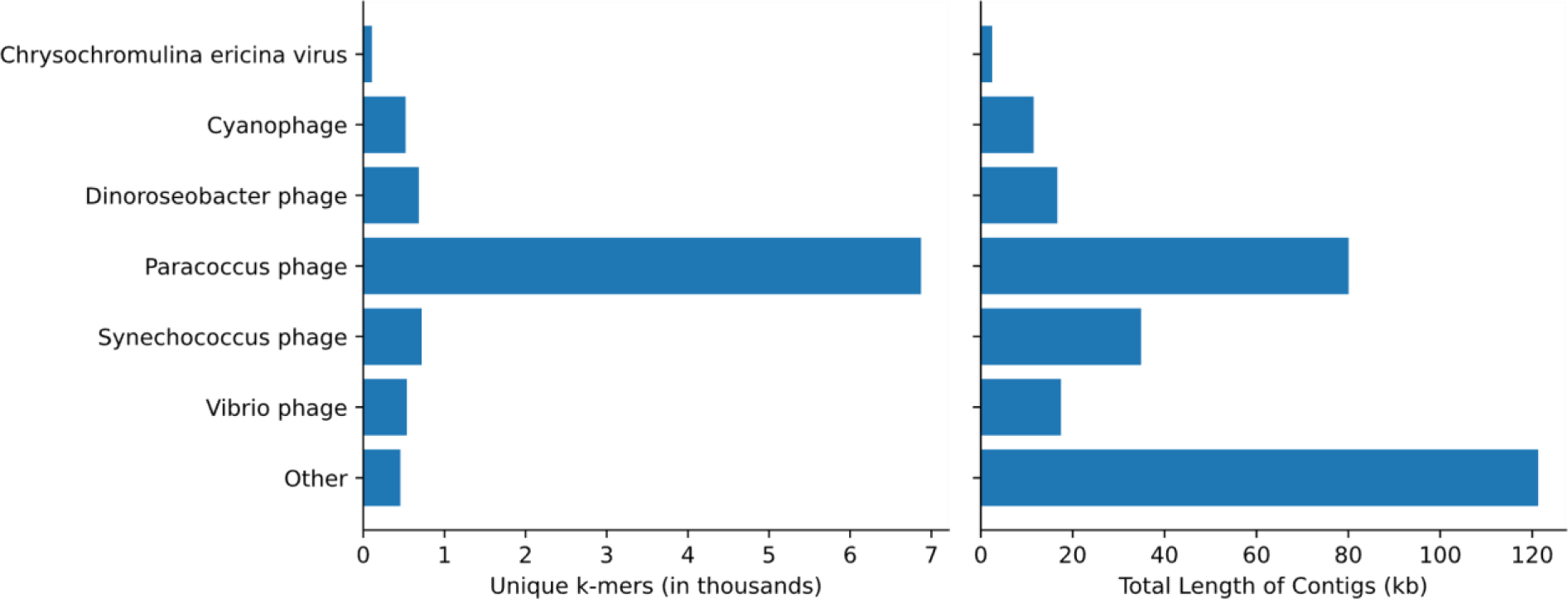
Classifications of assembled viral contigs showing the number of unique k-mers (in thousands) and the total sequence length (in kb) assembled for every genus.

When combining the filtered diseased reads from every coral species for assembly, the reads originating from *S. intersepta* samples represented a majority of the reads (89.8%). Therefore, we repeated the previous steps without *S. intersepta* reads and assembled the reads from the other three coral species. This resulted in 27,759 assembled contigs, of which 2,010 (7.2%) were classified by KrakenUniq, and only six matched viruses (five *Pseudoalteromonas* phages and one *Synechococcus* phage).

### Candidate genus (*Vibrio*) investigation

To explore the utility of our pipeline at a finer taxonomic level, we selected a candidate genus, *Vibrio*. We first investigated reads classified at the *Vibrio* species level. Vibrio reads were not able to be classified to the species level in 6.8%, 1.0%, 8.0%, and 11% of *S. intersepta, D. labyrinthiformis, D.stokesii*, and *M. meandrites*, respectively. The k-mers stemming from each *Vibrio* species for each host coral species pooled are displayed in Figure 5a and an overview of *Vibrio* read counts found in each colony sampled comprising the pooled coral species is shown in Figure 5b. *V. europaeus* and *V. tubiashii*, represented a large portion (37% combined) of the *Vibrio* k-mers in *S. intersepta* (Figure 5a). *V. mediterranei* dominated in *D. labyrinthiformis* (96%) and *D. stokesii* (67%) but was muted in the other coral species (Figure 5a); however, one colony per species appeared to be responsible for these high proportions: *D. labyrinthiformis* colony 57 and *D. stokesii* colony 63 (Figure 5b). *V. sp. THAF190c* was the predominant *Vibrio* species (17%) in *M. meandrites*. In general, other species like *V. coralliilyticus*, *V. harveyi*, *V. owensii*, and *V. sp. THAF100* appeared consistently in all coral species, but never in high proportions, but *V. tubiashii* and *V. owensii* contributed more than 5% of the reads in 10 samples each (Figure 5b).

**Figure 5.**
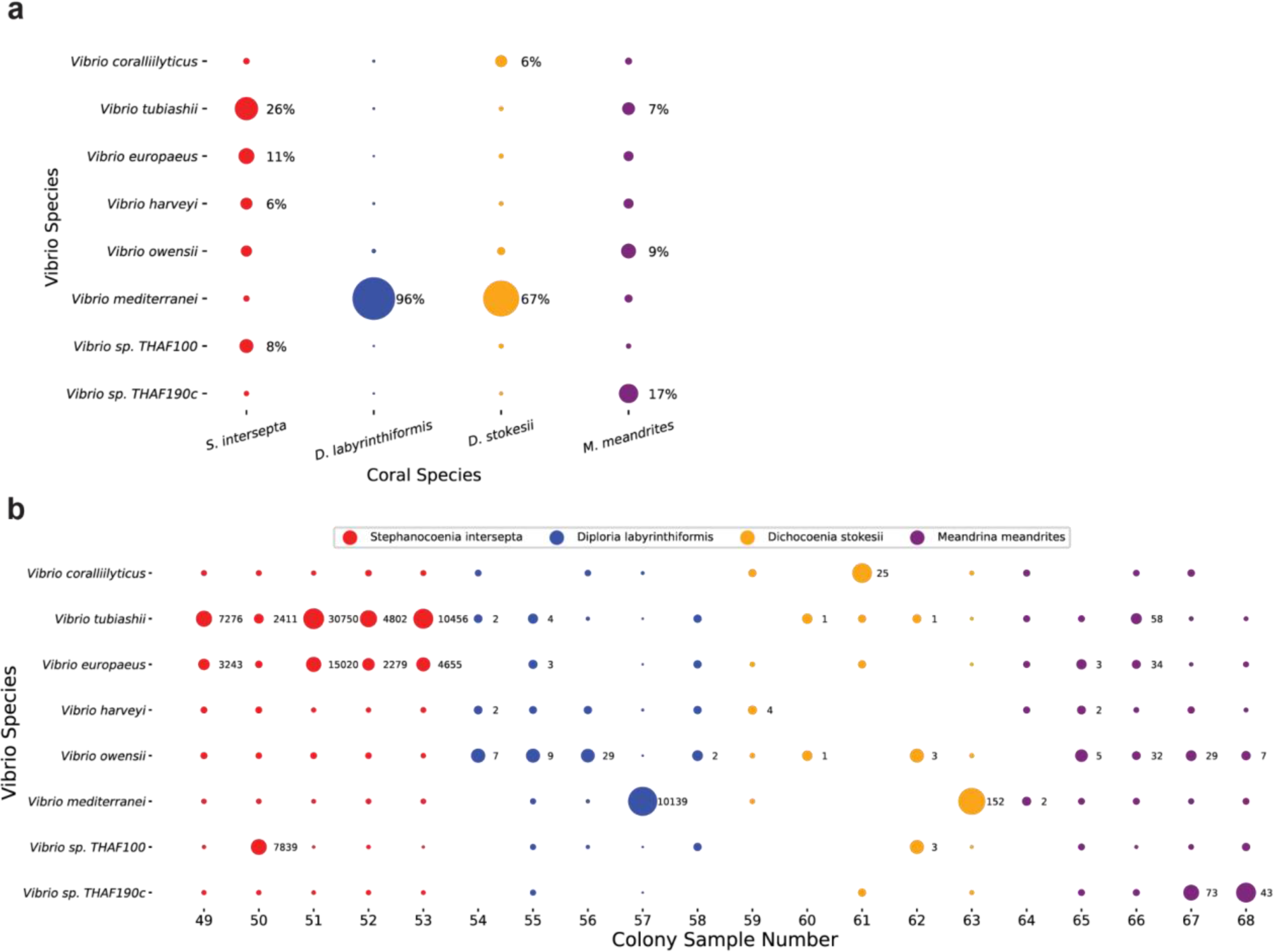
Species-level KrakenUniq classifications of *Vibrio* genus reads in SCTLD lesions. (**a**) The proportion of unique k-mers assigned to each *Vibrio* species grouped by samples from the same coral species. The size of the dots is relative to the proportion of unique k-mers assigned to the *Vibrio* species in each coral species. Those with over 5% are annotated with their proportion. (**b**) The species level read count proportions of the *Vibrio* for every sample. The size of the dots is relative to the proportion of read counts assigned to *Vibrio* species in each sample. The x-axis is labeled with the colony sample number, which corresponds to those assigned in the original analysis of this dataset (Rosales *et al*. 2020). Classifications that represent at least 5% of the *Vibrio* species in the sample are annotated with the number of reads.

Secondly, eight draft genomes from *V. coralliilyticus* strains isolated from a previous SCTLD study (Ushijima *et al*. 2020) allowed us to continue exploring the use of our pipeline and determine whether we identified the same *V. coralliilyticus* strains within our study. Reads classified by KrakenUniq as *Vibrio* in *S. intersepta* and *D. labyrinthiformis* were extracted and aligned to the Ushijima *et al*. (2020) draft genomes. Between 3.76% to 4.02% of the reads mapped to the *V. coralliilyticus* strains, while 7.54% of reads mapped to the McD22-P3 *Vibrio* strain, which was the control strain, and not a *V. coralliilyticus* strain (Supp. Table S4). As would be expected, the proportions of reads that are aligned are similar to the proportions of reads that were classified as *V. coralliilyticus* by KrakenUniq (Figure 5a).

## Discussion

In this study, we used previously published sequencing data from coral affected by SCTLD and developed a novel metagenomic analysis pipeline to explore the microbial communities present in those data. The data consisted of samples from four coral species collected from Florida’s coral reefs during a SCTLD outbreak. To investigate the microbial taxonomy of these samples, the previous study used small subunit rRNA gene assemblies and metagenome-assembled genomes (MAGs). Our investigation differed by using the Kraken software suite and focusing on unique k-mer count data to understand abundances. By predominantly working with k-mers instead of MAGs, we maximized the utility of the read data, as our approach allowed us to capture reads that may have been discarded during MAG assembly and binning. Samples with high intrapopulation diversity can provide challenges in MAG assembly and binning (Ramos-Barbero et al. 2019; Meziti et al. 2021), which may have hindered the generation of a MAG bin in the previous analysis. The previous work also did not filter out host sequences because reference genomes do not exist for the sampled coral species. However, here we applied a novel technique to filter these reads by using data derived from apparently healthy (AH) samples as a surrogate for reference genomes. This allowed us to examine unique sequences from DL samples by approximating a species-specific host coral genome and reduce computational load. In addition, in this study, we investigated the SCTLD DNA virome, which has not been previously reported.

The families *Rhodobacteraceae* and *Flavobacteriaceae* were found to be associated with SCTLD in our protein analysis, consistent with other SCTLD studies (reviewed in Papke *et al*. 2024), including previous examinations of the same sample set (Rosales *et al*. 2020, 2022). *Rhodobacteraceae* is one of the most common bacterial families associated with coral diseases (Gignoux-Wolfsohn *et al*. 2017) in diverse geographic locations, but no member has been identified as a causative coral disease agent (Mouchka *et al*. 2010), and members of this group are broadly found across ocean habitats and fulfill diverse ecological functions (Brinkhoff *et al*. 2008). *Flavobacteriaceae* has been enriched in White Band Disease in the Scleractinia staghorn coral *Acropora cervicornis* (Gignoux-Wolfsohn and Vollmer 2015), but has never been identified as a causative agent in coral tissue loss. In SCTLD, *Flavobacteriaceae* has previously been found enriched in unaffected (non-lesion) tissue from diseased colonies, potentially indicating colony stress and initial dysbiosis due to disease (Rosales *et al*. 2023). In contrast to the protein analysis, which used read count data, our DNA-level analysis using k-mers did not detect a singular genus belonging to *Rhodobacteraceae* or *Flavobacteriaceae* in high proportions among the unique diseased reads, providing support for the idea that members of these two families are likely a diverse set of bacteria species that associate with SCTLD opportunistically.

In our DNA-level analysis, *Vibrionaceae* and *Synechococcaceae* were among the most abundant families within the unique DL reads and these high abundances showcase how the k-mer based abundance approach can provide a different perspective of highly abundant taxa in particular samples. These families were not observed as highly abundant in the MMSeqs2 protein analysis; however, we were primarily interested in whether new candidates emerged from the MMSeqs2 analysis, not the relative proportions of the candidates. Interestingly, the genera *Synechococcus* and *Vibrio* were not detected in the previous analysis of these data (Rosales *et al*. 2022). In the 16S rRNA metabarcoding study of these same samples, *Synechococcus* and *Vibrio* were differentially abundant in *M. meandrites* samples (Rosales *et al*. 2020). These results did not show a clear trend as *Synechococcus* was present across multiple AH samples and *Vibrio* was not prevalent across DL coral samples. In this study, *Synechococcus* was the most or second most proportionally abundant genus across all four coral species. *Synechococcus* belong to phylum *Cyanobacteria*, which are photosynthetic picoplankton (Kim *et al*. 2018), and typically not involved in pathogenesis. Even so, *Synechococcus* have been enriched in other SCTLD studies that compared healthy colonies and healthy tissue on diseased colonies, and it was hypothesized that their increase in abundance is a response to disease stress (Rosales *et al*. 2023). The high proportion of *Synechococcus* in this study supports the suggestion that *Synechococcus* may have some role in microbial community interactions during SCTLD.

*Vibrio* also have been associated with SCTLD (reviewed in Papke *et al*. 2024), including in these samples (Rosales *et al*. 2020), but in contrast to *Synechococcus*, *Vibrio* have been associated with other coral tissue loss diseases, such as white pox disease (Kemp *et al*. 2018) and yellow band disease (Cervino *et al*. 2008), and diseases in other marine organisms, such as sponges (Dincturk *et al*. 2023) and crustaceans (de Souza Valente and Wan 2021). In three coral species analyzed here, *Vibrio* was one of the two most abundant genera, suggesting it may be associated with SCTLD tissue loss. In the fourth species, *M. meandrites, Vibrio* was only the sixth most abundant genus. However, *M. meandrites* had the fewest AH reads to create the database used for filtering the DL reads (Table 1), which may reduce the effectiveness of our novel pipeline that relies on reads from control groups. This was also a surprising result, since as mentioned, *Vibrio* were abundant in the 16S rRNA *M. meandrites* analysis of these samples (Rosales *et al*., 2020).

When using our pipeline to explore a species-level analysis within the candidate genus *Vibrio*, we found read matches to *V. mediterranei/shilonii* (Tarazona et al. 2014), *V. coralliilyticus*, *V. harveyi*, and *V. owensii* – all known coral pathogens associated with bleaching and tissue loss (Kushmaro *et al*. 1997; Ben-Haim *et al*. 2003; Luna *et al*. 2007; Ushijima *et al*. 2012; Munn 2015). *V. mediterranei/shilonii*, which comprised a majority of the classified *Vibrio* species in *D. labyrinthiformis* and *D. stokesii,* was previously found to be responsible for the annual bleaching of the scleractinian coral *Oculina patagonica* off the Israeli coast from 1993 (Kushmaro *et al*. 1996, 1997) to 2003 (Reshef *et al*. 2006). Additionally, when *V. mediterranei*/*shilonii* is experimentally introduced together with *V. coralliilyticus* to healthy corals, they appear to have synergistic pathogenic effects (Rubio-Portillo *et al*. 2014). Given these associations, *V. mediterranei*/*shilonii*, may be of particular interest in future SCTLD studies as potentially increasing the virulence of SCTLD, as does *V. coralliilyticus* (Ushijima *et al*. 2020), which also appeared in our samples. However, the very low similarity between the cultured *V. coralliilyticus* sequences in Ushijima *et al*. (2020) and our *Vibrio* sequences leads us to believe that multiple *Vibrio* species may be involved in SCTLD lesion development. It is important to note that our picture of the SCTLD microbiome is restricted by the genomes in the databases used. The *Vibrio* genus has been found to have a large degree of genetic plasticity between species (Gu et al. 2009), so while the classifications to different *Vibrio* species may truly represent the presence of an array of *Vibrio* species, it may instead be the result of various reads from a novel *Vibrio* species matching different *Vibrio* species based on closest genomic similarity. Therefore, while matches to different *Vibrio* species and their potential role in SCTLD may offer some insights, a more robust interpretation is to consider the implications of disease association by *Vibrio* at the genus level.

The etiology of SCTLD has been hypothesized to be viral (Work *et al*. 2021), and gene expression data show that there is an increase in coral viral immune response in corals with SCTLD (Beavers *et al*. 2023). Researchers have explored the potential role of RNA viruses in SCTLD, but no RNA viruses have been found exclusively in corals with SCTLD (Veglia *et al*. 2022) and these viruses are likely ubiquitous in corals without any potential relationship to SCTLD (Howe-Kerr *et al*. 2023). However, the involvement of DNA viruses has not previously been explored. Our data show the majority of DNA viruses in diseased samples represent phages. Not surprisingly, phage sequences correspond with some of the most abundant bacteria identified in this study, such as *Rhodobacteraceae, Vibrionaceae,* and *Synechococcaceae*. The *Paracoccus* phage, which infects *Rhodobacteraceae,* and the *Vibrio* phage would be interesting to further explore as potential avenues for disease mitigation. In addition to phages, sequences were found with similarities to the *Chrysochromulina ericina* virus. However, with only two contigs and little coverage of its genome, we do not believe this virus plays a role in SCTLD. Thus, we did not find any DNA viruses with a definitive association with SCTLD. Future studies may consider viral enrichment protocols prior to sequencing to help better characterize the SCTLD DNA virome.

In addition to differences in the bacterial and viral communities, members of *Symbiodiniaceae* were found to be abundant in DL samples, particularly in *M. meandrites*. SCTLD disrupts the relationship between the host coral and its *Symbiodiniaceae* through symbiont necrosis and peripheral nuclear chromatin condensation, among other physiological changes (Landsberg *et al*. 2020). This may result from an increase in *rab7* signaling among the *Symbiodiniaceae* to degrade dead and dysfunctional cells through endocytic phagosomes (Beavers *et al*. 2023). The *Symbiodiniaceae* DNA identified in diseased samples in this study may be a byproduct of this necrosis and degradation of the symbiont. This was especially notable in *M. meandrites,* which was the coral in this study most susceptible to acute tissue loss and mortality from SCTLD (Precht *et al*. 2016); this accelerated tissue loss may lead to higher levels of dead and dysfunctional symbionts being produced during visual lesion progression in *M. meandrites* than in other coral species.

Although, this pipeline was initially developed under the one-pathogen-one-disease assumption for humans, our results show that we can identify a consortium of putative pathogens with this method (i.e., *Rhodobacteraceae*, *Flavobacteriaceae*, and *Vibrio*). However, this pipeline is limited by the amount of prior information about pathogens in the species investigated; for example, human pathogens are well represented in metagenomic classification databases, whereas non-model organism pathogens may not be, which can lead to a higher rate of unclassified or misclassified reads. In addition, disease states are more clearly defined in humans than in non-model organisms, making the assumptions about sample health more reliable in human analyses.

## Conclusions

In all, we provide a novel pipeline to understand coral disease, and further investigate the role of bacteria and DNA viruses in SCTLD. Our new pipeline found both incongruities and parallels with the original analysis of these data. For instance, both studies revealed a prevalence of *Rhodobacteraceae* and *Flavobacteriaceae* in lesion samples. One of the major differences between the studies was the dominance of *Synechococcus* and *Vibrio* in this study, which was likely driven by the use of k-mers to quantify taxonomic abundance (as the protein analysis with read counts aligned more with previous results). In addition to providing novel insights, our pipeline also reduced the computational load for downstream analysis, making large metagenomic analysis more accessible to those with less computational resources. This method is useful not only for coral scientists, but also for fields that study non-model organisms and lack comprehensive genomic resources. In corals, the lack of such resources can impede the progress of metagenomic analysis and the identification of potential pathogens. Specifically, host-associated metagenomes without a reference genome can dominate and confound metagenomic results (Rosales *et al*. 2022). As genome assemblies are unlikely to be available anytime soon for the over 30 species of coral affected by SCTLD, this pipeline offers a useful tool for the simultaneous examination of viruses, bacteria, and eukaryotic species living on or within the host tissue that may be associated with infection for any coral species or non-model organism.

## Data availability

The samples investigated in this study were downloaded from NCBI BioProject PRJNA576217. Supplemental materials are available at FigShare. Table S1 contains the average k-mers per read for every genus across all coral species. Table S2 contains the *Symbiodinaceae* protein classifications. Table S3 contains the virulence factors identified in the *Vibrio* contigs. Table S4 contains the alignment rates of reads classified as *Vibrio* to eight draft assemblies of *Vibrio* species previously isolated from SCTLD infected corals. The IDs of the unique diseased reads remaining after filtering for each coral species and the assembled contigs are also available at FigShare. The report files from the KrakenUniq and MMseqs2 analyses have been made available at the following repository: https://github.com/jheinz27/coral_results/tree/main.

## Acknowledgements

We thank Dr. Martin Steinegger and Dr. Christopher Pockrandt for their helpful advice and exploratory work on pathogen identification in Mariana crows, which helped inform the methods development here. We also thank Dr. Alaina Shumate for her valuable advice and guidance. The samples were collected under permits #FKNMS-2018-057 and #FKNMS-2017-100.

## Literature Cited

Aeby, G. S., B. Ushijima, J. E. Campbell, S. Jones, G. J. Williams et al., 2019 Pathogenesis of a tissue loss disease affecting multiple species of corals along the Florida reef tract. Front. Mar. Sci. 6.: 10.3389/fmars.2019.00678

Aiewsakun, P., E. M. Adriaenssens, R. Lavigne, A. M. Kropinski, and P. Simmonds, 2018 Evaluation of the genomic diversity of viruses infecting bacteria, archaea and eukaryotes using a common bioinformatic platform: steps towards a unified taxonomy. J. Gen. Virol. 99: 1331–1343. 10.1099/jgv.0.001110

Altschul, S. F., W. Gish, W. Miller, E. W. Myers, and D. J. Lipman, 1990 Basic local alignment search tool. J. Mol. Biol. 215: 403–410. 10.1016/S0022-2836(05)80360-2

Alvarez-Filip, L., F. J. González-Barrios, E. Pérez-Cervantes, A. Molina-Hernández, and N. Estrada-Saldívar, 2022 Stony coral tissue loss disease decimated Caribbean coral populations and reshaped reef functionality. Commun Biol 5: 440. 10.1038/s42003-022-03398-6

Amos, B., C. Aurrecoechea, M. Barba, A. Barreto, E. Y. Basenko et al., 2022 VEuPathDB: the eukaryotic pathogen, vector and host bioinformatics resource center. Nucleic Acids Res. 50: D898–D911. 10.1093/nar/gkab929

Beavers, K. M., E. W. Van Buren, A. M. Rossin, M. A. Emery, A. J. Veglia et al., 2023 Stony coral tissue loss disease induces transcriptional signatures of in situ degradation of dysfunctional Symbiodiniaceae. Nat. Commun. 14: 2915. 10.1038/s41467-023-38612-4

Ben-Haim, Y., F. L. Thompson, C. C. Thompson, M. C. Cnockaert, B. Hoste et al., 2003 Vibrio coralliilyticus sp. nov., a temperature-dependent pathogen of the coral Pocillopora damicornis. Int. J. Syst. Evol. Microbiol. 53: 309–315. 10.1099/ijs.0.02402-0

Benson, D. A., M. Cavanaugh, K. Clark, I. Karsch-Mizrachi, D. J. Lipman et al., 2017 GenBank. Nucleic Acids Res. 45: D37–D42. 10.1093/nar/gkw1070

Bourne, D. G., M. Garren, T. M. Work, E. Rosenberg, G. W. Smith et al., 2009 Microbial disease and the coral holobiont. Trends Microbiol. 17: 554–562. 10.1016/j.tim.2009.09.004

Breitwieser, F. P., D. N. Baker, and S. L. Salzberg, 2018 KrakenUniq: confident and fast metagenomics classification using unique k-mer counts. Genome Biol. 19: 198. 10.1186/s13059-018-1568-0

Breitwieser, F. P., and S. L. Salzberg, 2020 Pavian: interactive analysis of metagenomics data for microbiome studies and pathogen identification. Bioinformatics 36: 1303–1304. 10.1093/bioinformatics/btz715

Brinkhoff, T., H.-A. Giebel, and M. Simon, 2008 Diversity, ecology, and genomics of the Roseobacter clade: a short overview. Arch. Microbiol. 189: 531–539. 10.1007/s00203-008-0353-y

Callahan, B. J., P. J. McMurdie, M. J. Rosen, A. W. Han, A. J. A. Johnson et al., 2016 DADA2: High-resolution sample inference from Illumina amplicon data. Nat. Methods 13: 581–583. 10.1038/nmeth.3869

Cervino, J. M., F. L. Thompson, B. Gomez-Gil, E. A. Lorence, T. J. Goreau et al., 2008 The Vibrio core group induces yellow band disease in Caribbean and Indo-Pacific reef-building corals. J. Appl. Microbiol. 105: 1658–1671. 10.1111/j.1365-2672.2008.03871.x

Clark, A. S., S. D. Williams, K. Maxwell, S. M. Rosales, L. K. Huebner et al., 2021 Characterization of the Microbiome of Corals with Stony Coral Tissue Loss Disease along Florida’s Coral Reef. Microorganisms 9.: 10.3390/microorganisms9112181

de Souza Valente, C., and A. H. L. Wan, 2021 Vibrio and major commercially important vibriosis diseases in decapod crustaceans. J. Invertebr. Pathol. 181: 107527. 10.1016/j.jip.2020.107527

Dinçtürk, E., F. Öndes, L. Leria, and M. Maldonado, 2023 Mass mortality of the keratose sponge Sarcotragus foetidus in the Aegean Sea (Eastern Mediterranean) correlates with proliferation of Vibrio bacteria in the tissues. Front. Microbiol. 14: 1272733. 10.3389/fmicb.2023.1272733

Eberhart, C., Li, Z., Breitwieser, F. P., Lu, J., Jun, A. S. et al., 2017 Diagnosing corneal infections in formalin fixed specimens using next generation sequencing. Investigative Ophthalmology & Visual Science.

Gallot-Lavallée, L., G. Blanc, and J.-M. Claverie, 2017 Comparative Genomics of Chrysochromulina Ericina Virus and Other Microalga-Infecting Large DNA Viruses Highlights Their Intricate Evolutionary Relationship with the Established Mimiviridae Family. J. Virol. 91.: 10.1128/JVI.00230-17

Gallot-Lavallée, L., A. Pagarete, M. Legendre, S. Santini, R.-A. Sandaa et al., 2015 The 474-Kilobase-Pair Complete Genome Sequence of CeV-01B, a Virus Infecting Haptolina (Chrysochromulina) ericina (Prymnesiophyceae). Genome Announc. 3.: 10.1128/genomeA.01413-15

Gavriilidou, A., J. Gutleben, D. Versluis, F. Forgiarini, M. W. J. van Passel et al., 2020 Comparative genomic analysis of Flavobacteriaceae: insights into carbohydrate metabolism, gliding motility and secondary metabolite biosynthesis. BMC Genomics 21: 569. 10.1186/s12864-020-06971-7

Gignoux-Wolfsohn, S. A., F. M. Aronson, and S. V. Vollmer, 2017 Complex interactions between potentially pathogenic, opportunistic, and resident bacteria emerge during infection on a reef-building coral. FEMS Microbiol. Ecol. 93.: 10.1093/femsec/fix080.

Gignoux-Wolfsohn, S. A., and S. V. Vollmer, 2015 Identification of Candidate Coral Pathogens on White Band Disease-Infected Staghorn Coral. PLoS One 10: e0134416. 10.1371/journal.pone.0134416

Gihawi, A., Y. Ge, J. Lu, D. Puiu, A. Xu et al., 2023 Major data analysis errors invalidate cancer microbiome findings. MBio 14: e0160723. e0160723. 10.1128/mbio.01607-23

Göker, M., 2022 Filling the gaps: missing taxon names at the ranks of class, order and family. Int. J. Syst. Evol. Microbiol. 72.: 10.1099/ijsem.0.005638

Gu, J., J. Neary, H. Cai, A. Moshfeghian, S. A. Rodriguez et al., 2009 Genomic and systems evolution in Vibrionaceae species. BMC Genomics 10 Suppl 1: S11. 10.1186/1471-2164-10-S1-S11

Howe-Kerr, L. I., A. M. Knochel, M. D. Meyer, J. A. Sims, C. E. Karrick et al., 2023 Filamentous virus-like particles are present in coral dinoflagellates across genera and ocean basins. ISME J. 10.1038/s41396-023-01526-6

Huang, H., P. B. McGarvey, B. E. Suzek, R. Mazumder, J. Zhang et al., 2011 A comprehensive protein-centric ID mapping service for molecular data integration. Bioinformatics 27: 1190–1191. 10.1093/bioinformatics/btr101

Huntley, N., M. E. Brandt, C. C. Becker, C. A. Miller, S. S. Meiling et al., 2022 Experimental transmission of Stony Coral Tissue Loss Disease results in differential microbial responses within coral mucus and tissue. ISME Communications 2: 1–11. 10.1038/s43705-022-00126-3

Kim, Y., J. Jeon, M. S. Kwak, G. H. Kim, I. Koh et al., 2018 Photosynthetic functions of Synechococcus in the ocean microbiomes of diverse salinity and seasons. PLoS One 13: e0190266. 10.1371/journal.pone.0190266

Kemp, K. M., J. R. Westrich, M. S. Alabady, M. L. Edwards, and E. K. Lipp, 2018 Abundance and Multilocus Sequence Analysis of Vibrio Bacteria Associated with Diseased Elkhorn Coral (Acropora palmata) of the Florida Keys. Appl. Environ. Microbiol. 84.: 10.1128/AEM.01035-17

Kostic, A. D., D. Gevers, C. S. Pedamallu, M. Michaud, F. Duke et al., 2012 Genomic analysis identifies association of Fusobacterium with colorectal carcinoma. Genome Res. 22: 292–298. 10.1101/gr.126573.111

Kushmaro, A., Y. Loya, M. Fine, and E. Rosenberg, 1996 Bacterial infection and coral bleaching. Nature 380: 396–396. 10.1038/380396a0

Kushmaro, A., E. Rosenberg, M. Fine, and Y. Loya, 1997 Bleaching of the coral Oculina patagonica by Vibrio AK-1. Mar. Ecol. Prog. Ser. 147: 159–165. 10.3354/meps147159

Landsberg, J. H., Y. Kiryu, E. C. Peters, P. W. Wilson, N. Perry et al., 2020 Stony coral tissue loss disease in Florida is associated with disruption of host–zooxanthellae physiology. Front. Mar. Sci. 7.: 10.3389/fmars.2020.576013

Langmead, B., and S. L. Salzberg, 2012 Fast gapped-read alignment with Bowtie 2. Nat. Methods 9: 357–359. 10.1038/nmeth.1923

Li, D., R. Luo, C.-M. Liu, C.-M. Leung, H.-F. Ting et al., 2016 MEGAHIT v1.0: A fast and scalable metagenome assembler driven by advanced methodologies and community practices. Methods 102: 3–11. 10.1016/j.ymeth.2016.02.020

Lin, H., M. Yu, X. Wang, and X.-H. Zhang, 2018 Comparative genomic analysis reveals the evolution and environmental adaptation strategies of vibrios. BMC Genomics 19: 135. 10.1186/s12864-018-4531-2

Liu, B., D. Zheng, S. Zhou, L. Chen, and J. Yang, 2022 VFDB 2022: a general classification scheme for bacterial virulence factors. Nucleic Acids Res. 50: D912–D917. 10.1093/nar/gkab1107

Lu, J., N. Rincon, D. E. Wood, F. P. Breitwieser, C. Pockrandt et al., 2022 Metagenome analysis using the Kraken software suite. Nat. Protoc. 17: 2815–2839. 10.1038/s41596-022-00738-y

Lu, J., and S. L. Salzberg, 2018 Removing contaminants from databases of draft genomes. PLoS Comput. Biol. 14: e1006277. 10.1371/journal.pcbi.1006277

Luna, G. M., F. Biavasco, and R. Danovaro, 2007 Bacteria associated with the rapid tissue necrosis of stony corals. Environ. Microbiol. 9: 1851–1857. 10.1111/j.1462-2920.2007.01287.x

Meziti, A., L. M. Rodriguez-R, J. K. Hatt, A. Peña-Gonzalez, K. Levy et al., 2021 The Reliability of Metagenome-Assembled Genomes (MAGs) in Representing Natural Populations: Insights from Comparing MAGs against Isolate Genomes Derived from the Same Fecal Sample. Appl. Environ. Microbiol. 87. 10.1128/AEM.02593-20

Meyer, J. L., J. Castellanos-Gell, G. S. Aeby, C. C. Häse, B. Ushijima et al., 2019 Microbial Community Shifts Associated With the Ongoing Stony Coral Tissue Loss Disease Outbreak on the Florida Reef Tract. Front. Microbiol. 10: 2244. 10.3389/fmicb.2019.02244

Mouchka, M. E., I. Hewson, and C. D. Harvell, 2010 Coral-associated bacterial assemblages: current knowledge and the potential for climate-driven impacts. Integr. Comp. Biol. 50: 662–674. 10.1093/icb/icq061

Munn, C. B., 2015 The Role of Vibrios in Diseases of Corals. Microbiol Spectr 3.: 10.1128/microbiolspec.VE-0006-2014

National Center for Biotechnology Information (NCBI) Scleractinia [Internet]. Bethesda (MD): National Library of Medicine (US), National Center for Biotechnology Information; [1988] [cited 2023 Nov 26]. Available from: https://www.ncbi.nlm.nih.gov/datasets/taxonomy/6125/

Neely, K. L., K. A. Macaulay, E. K. Hower, and M. A. Dobler, 2020 Effectiveness of topical antibiotics in treating corals affected by Stony Coral Tissue Loss Disease. PeerJ 8: e9289. 10.7717/peerj.9289

Papke, E., A. Carreiro, C. Dennison, J. M. Deutsch, L. M. Isma et al., 2024 Stony coral tissue loss disease: a review of emergence, impacts, etiology, diagnostics, and intervention. Front. Mar. Sci. 10.: 10.3389/fmars.2023.1321271

Precht, W. F., B. E. Gintert, M. L. Robbart, R. Fura, and R. van Woesik, 2016 Unprecedented Disease-Related Coral Mortality in Southeastern Florida. Sci. Rep. 6: 31374. 10.1038/srep31374

Quast, C., E. Pruesse, P. Yilmaz, J. Gerken, T. Schweer et al., 2013 The SILVA ribosomal RNA gene database project: improved data processing and web-based tools. Nucleic Acids Res. 41: D590–6. 10.1093/nar/gks1219

Reshef, L., O. Koren, Y. Loya, I. Zilber-Rosenberg, and E. Rosenberg, 2006 The coral probiotic hypothesis. Environ. Microbiol. 8: 2068–2073. 10.1111/j.1462-2920.2006.01148.x

Ramos-Barbero, M. D., A.-B. Martin-Cuadrado, T. Viver, F. Santos, M. Martinez-Garcia et al., 2019 Recovering microbial genomes from metagenomes in hypersaline environments: The Good, the Bad and the Ugly. Syst. Appl. Microbiol. 42: 30–40. 10.1016/j.syapm.2018.11.001

Rosales, S. M., A. S. Clark, L. K. Huebner, R. R. Ruzicka, and E. M. Muller, 2020 Rhodobacterales and Rhizobiales Are Associated With Stony Coral Tissue Loss Disease and Its Suspected Sources of Transmission. Front. Microbiol. 11: 681. 10.3389/fmicb.2020.00681

Rosales, S. M., L. K. Huebner, A. S. Clark, R. McMinds, R. R. Ruzicka et al., 2022 Bacterial Metabolic Potential and Micro-Eukaryotes Enriched in Stony Coral Tissue Loss Disease Lesions. Frontiers in Marine Science 8.: 10.3389/fmars.2021.776859

Rosales, S. M., L. K. Huebner, J. S. Evans, A. Apprill, A. C. Baker et al., 2023 A meta-analysis of the stony coral tissue loss disease microbiome finds key bacteria in unaffected and lesion tissue in diseased colonies. ISME Commun 3: 19. 10.1038/s43705-023-00220-0

Rubio-Portillo, E., P. Yarza, C. Peñalver, A. A. Ramos-Esplá, and J. Antón, 2014 New insights into Oculina patagonica coral diseases and their associated Vibrio spp. communities. ISME J. 8: 1794–1807. 10.1038/ismej.2014.33

Salzberg, S. L., F. P. Breitwieser, A. Kumar, H. Hao, P. Burger et al., 2016 Next-generation sequencing in neuropathologic diagnosis of infections of the nervous system. Neurol Neuroimmunol Neuroinflamm 3: e251. 10.1212/NXI.0000000000000251

Schmieder, R., and R. Edwards, 2011 Fast identification and removal of sequence contamination from genomic and metagenomic datasets. PLoS One 6: e17288. 10.1371/journal.pone.0017288

Shilling, E. N., I. R. Combs, and J. D. Voss, 2021 Assessing the effectiveness of two intervention methods for stony coral tissue loss disease on Montastraea cavernosa. Sci. Rep. 11: 8566. 10.1038/s41598-021-86926-4

Steinegger, M., and J. Söding, 2017 MMseqs2 enables sensitive protein sequence searching for the analysis of massive data sets. Nat. Biotechnol. 35: 1026–1028. 10.1038/nbt.3988

Studivan, M. S., R. J. Eckert, E. Shilling, N. Soderberg, I. C. Enochs et al., 2023 Stony coral tissue loss disease intervention with amoxicillin leads to a reversal of disease-modulated gene expression pathways. Mol. Ecol. 32: 5394–5413. 10.1111/mec.17110

Suzek, B. E., Y. Wang, H. Huang, P. B. McGarvey, C. H. Wu et al., 2015 UniRef clusters: a comprehensive and scalable alternative for improving sequence similarity searches. Bioinformatics 31: 926–932. 10.1093/bioinformatics/btu739

Tarazona, E., T. Lucena, D. R. Arahal, M. C. Macián, M. A. Ruvira et al., 2014 Multilocus sequence analysis of putative Vibrio mediterranei strains and description of Vibrio thalassae sp. nov. Syst. Appl. Microbiol. 37: 320–328. 10.1016/j.syapm.2014.05.005

UniProt: the universal protein knowledgebase in 2021, 2021 Nucleic Acids Res. 49: D480–D489. 10.1093/nar/gkaa1100

Ushijima, B., A. Smith, G. S. Aeby, and S. M. Callahan, 2012 Vibrio owensii induces the tissue loss disease Montipora white syndrome in the Hawaiian reef coral Montipora capitata. PLoS One 7: e46717. 10.1371/journal.pone.0046717

Ushijima, B., J. L. Meyer, S. Thompson, K. Pitts, M. F. Marusich et al., 2020 Disease Diagnostics and Potential Coinfections by Vibrio coralliilyticus During an Ongoing Coral Disease Outbreak in Florida. Front. Microbiol. 11: 569354. 10.3389/fmicb.2020.569354

Veglia, A. J., K. Beavers, E. W. Van Buren, S. S. Meiling, E. M. Muller et al., 2022 Alphaflexivirus Genomes in Stony Coral Tissue Loss Disease-Affected, Disease-Exposed, and Disease-Unexposed Coral Colonies in the U.S. Virgin Islands. Microbiol Resour Announc 11: e0119921. 10.1128/mra.01199-21

Walton, C. J., N. K. Hayes, and D. S. Gilliam, 2018 Impacts of a regional, multi-year, multi-species coral disease outbreak in southeast Florida. Front. Mar. Sci. 5.: 10.3389/fmars.2018.00323

Wilson, M. R., H. A. Sample, K. C. Zorn, S. Arevalo, G. Yu et al., 2019 Clinical Metagenomic Sequencing for Diagnosis of Meningitis and Encephalitis. N. Engl. J. Med. 380: 2327–2340. 10.1056/NEJMoa1803396

Work, T. M., T. M. Weatherby, J. H. Landsberg, Y. Kiryu, S. M. Cook et al., 2021 Viral-like particles are associated with Endosymbiont pathology in Florida corals affected by stony coral tissue loss disease. Front. Mar. Sci. 8.: 10.3389/fmars.2021.750658

