## Supplemental Figures and Methods for "Novel metagenomics analysis of stony coral tissue loss disease"

### Supplemental Materials

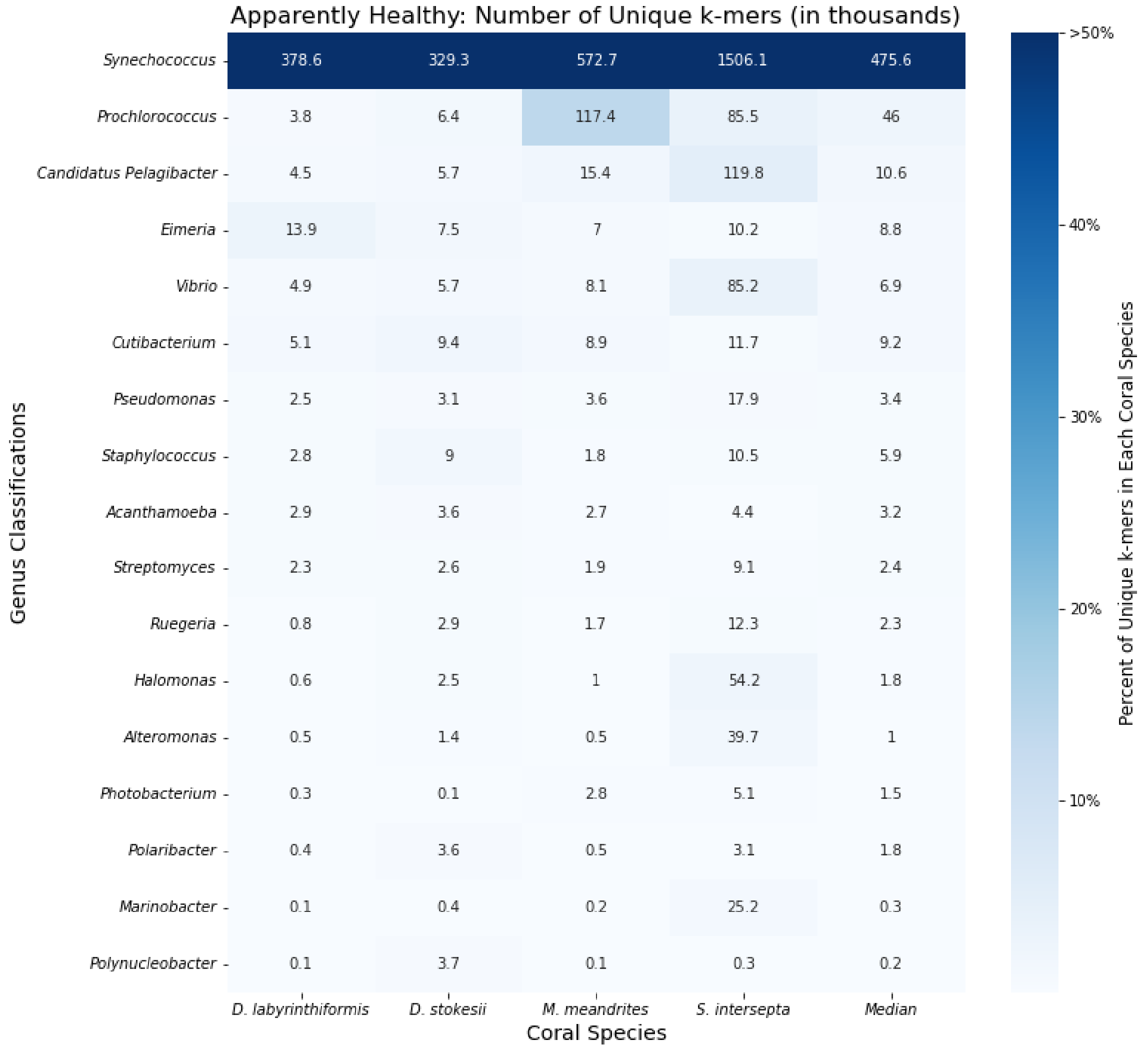


**Figure S1:** Unique k-mer counts from KrakenUniq genus-level classifications of Apparently Healthy (AH) sample reads for every coral species. The intensity of the shading represents the percent of total unique k-mers assigned to the genus. Each box is annotated with the number of unique k-mers (in thousands) assigned to the genus.

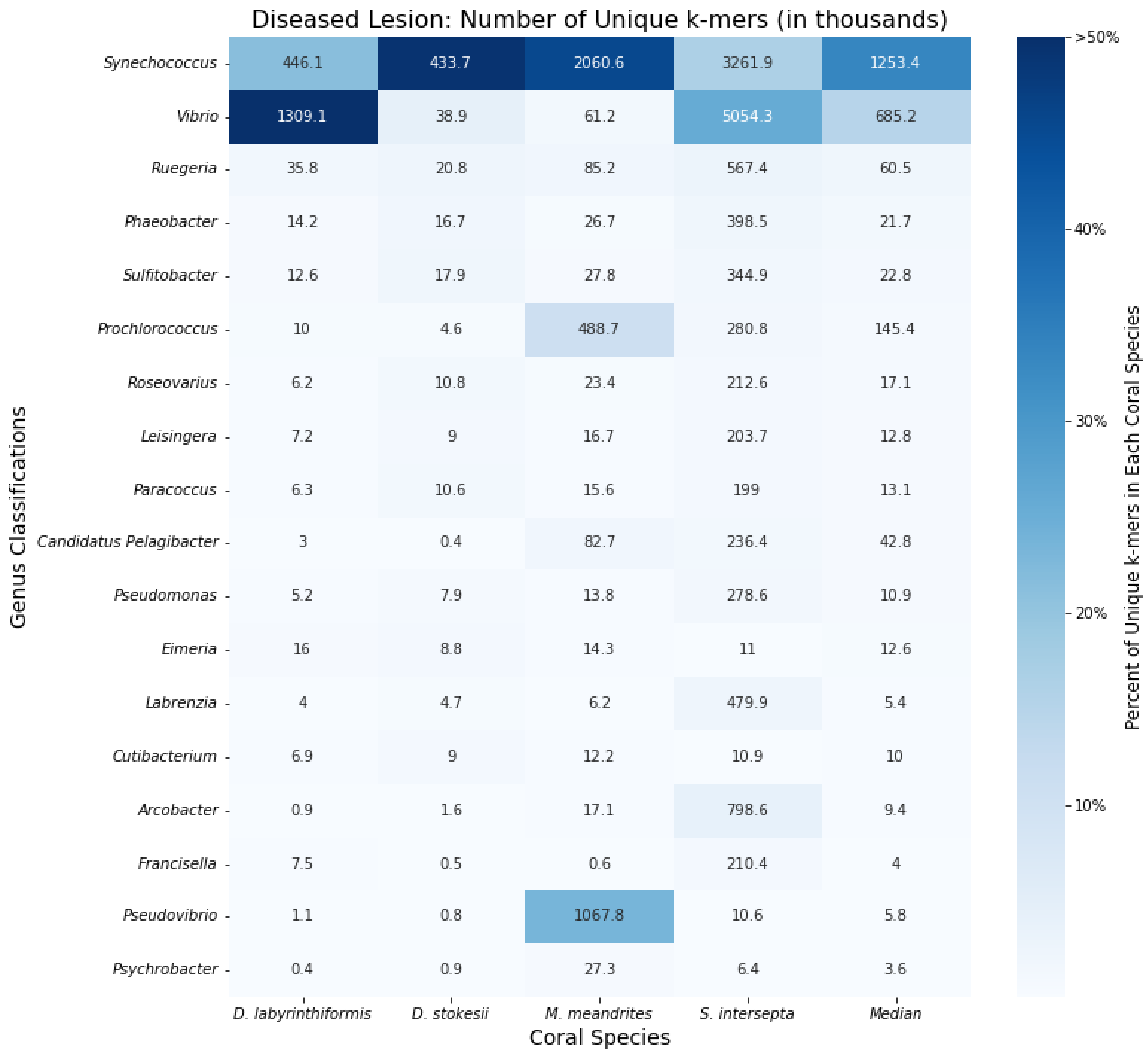


**Figure S2:** Unique k-mer counts from KrakenUniq genus-level classifications of all unfiltered Diseased Lesion (DL) sample reads for every coral species. The intensity of the shading represents the percent of total unique k-mers assigned to the genus. Each box is annotated with the number of unique k-mers (in thousands) assigned to the genus.

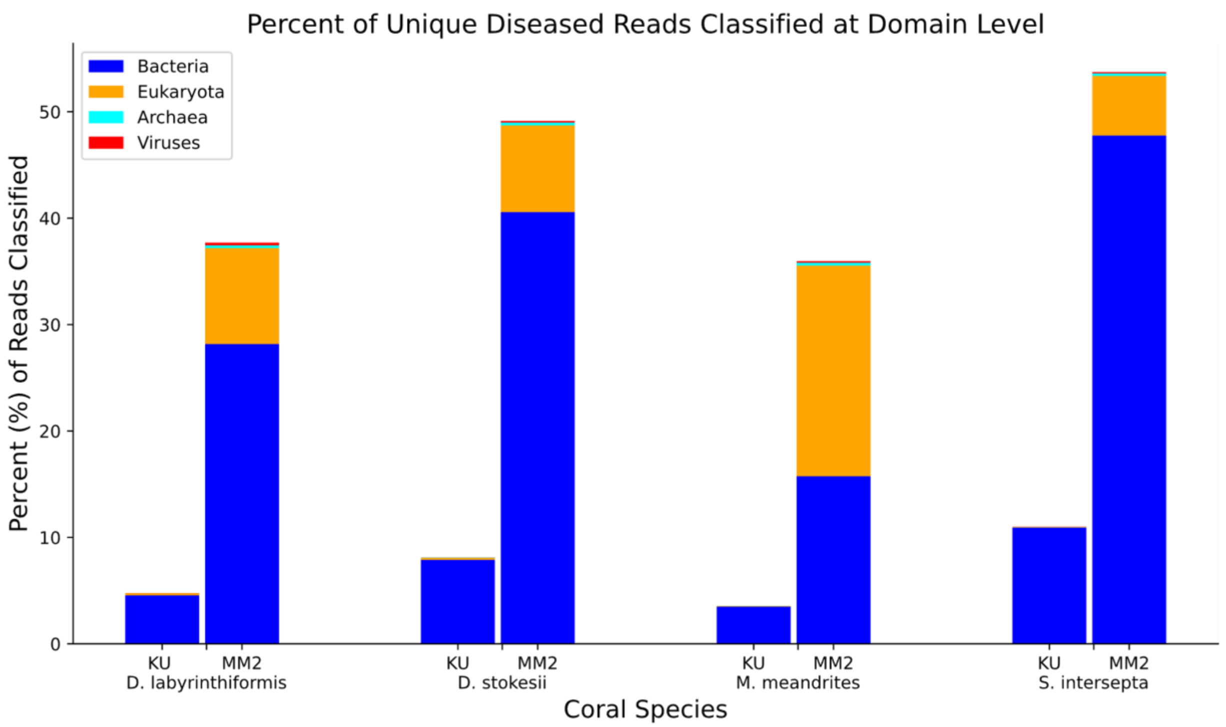


**Figure S3** Proportions of unique diseased reads classified by KrakenUniq (KU) and MMseqs2 (MM2), subdivided by superkingdom, for each coral species.

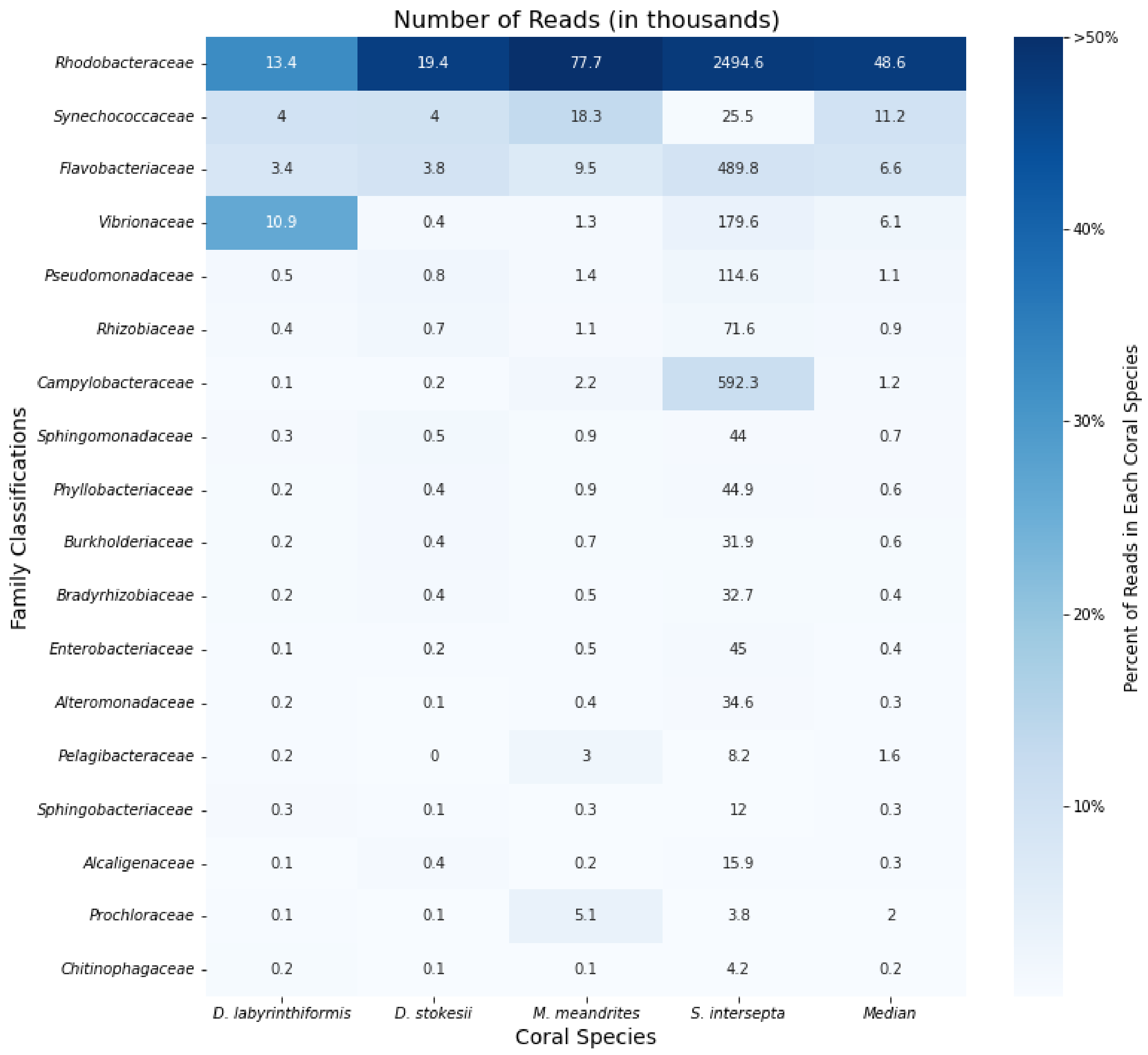


**Figure S4:** Read count from KrakenUniq family-level classifications of unique diseased reads for every coral species. The intensity of the shading represents the percent of total reads assigned to the family. Each box is annotated with the number of reads (in thousands) assigned to the genus.

Table S1- Average k-mers per read for every genus across all coral species.

Table S2- *Symbiodiniaceae* proteins that had at least one unique diseased read align to the UniRef50 cluster (paired reads classified separately) for each coral species.

Table S3- Hits to the Virulence Factor Database by assembled contigs that were classified as *Vibrio* by KrakenUniq.

Table S4- Alignment rates of reads classified as *Vibrio* to eight draft assemblies of *Vibrio* species isolated from SCTLD infected corals by Ushijima *et al.* 2020

### Supplemental Methods

#### Filtering reads with a healthy coral reference database

A database was for each coral species was created with the command:

```
krakenuniq-build –db pooled_healthy_reads --threads 32 --kmer-len 29
```

Each read k-mer found in the healthy reads was stored under the same taxonomy ID. The diseased lesion reads from the same species were then classified using the created database with the following command:

```
krakenuniq --db database_from_healthy_reads --threads 32 --paired --report
diseased_against_healthy.kuniqreport pooled_1.fasta pooled_2.fasta >
diseased_against_healthy.kuniq
```

Only unclassified reads were kept for downstream analysis.

#### KrakenUniq read classification

This command generated report files which list the number of reads and unique k-mers for all taxa, including counts at the strain, species, genus, and higher levels.

```
krakenuniq --db /ccb/salz8-4/data/krakendbs/krakendb-2020-08-16-
all_pluseupath/ --threads 16 --paired --report results.kuniqreport
filtered_reads_1.fasta filtered_reads_2.fasta > results.kuniq
```

#### MMseqs2 read classification

The flowing commands were used to clasify the paired read files individually with the Mmseq2 easy-taxonomy workflow against the UniRef50 database.

```
mmseqs easy-taxonomy filtered_reads_1.fasta uniref50_db
filtered_reads_results_1 tmp --threads 16
```

```
mmseqs easy-taxonomy filtered_reads_1.fasta uniref50_db
filtered_reads_results_1 tmp --threads 16
```

The report files were sorted by most abundant families using the command:

```
awk -F'\t' '$4 == "family" {print $0}' results_report | sort -nrk2,2
```
